# Influence of age and gender on taste function of healthy subjects

**DOI:** 10.1101/2019.12.12.873976

**Authors:** Jing-Jie Wang, Kai-Li Liang, Wen-Jiun Lin, Chih-Yi Chen, Rong-San Jiang

**Affiliations:** Institute of Medicine, Chung Shan Medical University, Taichung, Taiwan; Department of Otolaryngology, Taichung Veterans General Hospital, Taichung, Taiwan; Faculty of Medicine, National Yang-Ming Medical University, Taipei, Taiwan; School of Medicine, Chung Shan Medical University, Taichung, Taiwan; Division of Thoracic Surgery, Chung Shan Medical University Hospital, Taichung, Taiwan; Department of Medical Research, Taichung Veterans General Hospital, Taichung, Taiwan; Rong Hsing Research Center for Translational Medicine, National Chung Hsing University, Taichung, Taiwan

## Abstract

**Objectives:** To determine the influences of age and gender on the taste functions of healthy Taiwanese.

**Methods:** We evaluated taste functions of healthy Taiwanese using the whole mouth suprathreshold taste test, along with the taste quad test. In the whole-mouth test, we applied in a counterbalanced order sweet, sour, salty, and bitter solutions, each at 5 different suprathreshold concentrations to subjects, who were instructed to sip and swish in mouth twice. Each subject had to indicate the taste quality, and to rate the intensity and unpleasantness/pleasantness of each taste of the solutions. In the quad test, the 4 quadrants of the tongue surface were tested by dripping one concentration of sweet, sour, salty, or bitter solutions for 6 times. Subjects then indicated the taste quality, and rated the intensity of the solution.

**Results:** Subjects were divided into groups based on their gender and age: 20-39 years, 40-59 years, or ≥ 60 years. We found that in the whole mouth taste test, the total correct identification score dropped with age. But identifying sweet and salty qualities was not affected by age. No differences were found between male and female, except women scored better than men for sweet quality in the age group of 40-59 years. The total correct identification score of the taste quad test also decreased with an increased age of the taste quad test, without gender differences.

**Conclusion:** Both age and gender affected the taste functions in healthy Taiwanese to some extent, and differences were dependent on age, tongue region, and taste quality.

## Introduction

Taste dysfunction was estimated in prevalence to affect 26.3 million people in the US according to a 2016 nationwide survey [1]. It is generally accepted that our smell ability declines as we age [2]. However, the effect of aging on taste function is considered small and varies according to individuals [3]. A systemic review of aging effects on taste function reported that taste perception declines with aging, although the extents of decline differ across studies [4]. It is also known that gender affects taste preference, detection threshold, and reactivity to taste stimuli [5].But the exact nature of such gender effect remains illusive [5]. Gudziol and Hummel [6] used the ‘three-drop test’ to study taste function in a population of Europeans, and found that women have taste functions more sensitive than men. Another study also reported that gender affects the perception of sour and bitter tastes [7].

Taste function is rarely investigated in Asian populations. Yong et. al., [8] investigated the effect of age and gender on taste function in 90 healthy Chinese adults, using the same method of Gudziol and Hummel [6]. They found no effects of age or gender on taste function. They proposed that eating habits may influence the taste results, and that Asians are more sensitive to tastants. However, in their study, only subjects <65 years were analyzed.

Taste function has been determined using both chemical and electrical stimuli. Several methods have been developed to present chemical stimuli to human subjects, including ‘sipping & spitting’, tastant strips, taste tablets, cotton swabs, and discs [9-11]. Solution-based taste tests have known reliability [6]. In general, taste tests are either the whole-mouth test or the regional test [12]. The whole-mouth test provides general taste sensitivity [13], while the regional test can detect gustatory blind regions on the tongue [14]. In order to further clarify the influence of age and gender on taste functions, specifically in Asian populations, we conducted here a study to investigate the taste function in healthy Taiwanese, using the solution-based taste tests (same as those used at the Smell & Taste Center of the University of Pennsylvania).

## Materials and methods

### Ethical statements

This study was approved by the Institutional Review Board of Taichung Veterans General Hospital, Taiwan (IRB number: CF18048A). Informed written consents were collected from all enrolled subjects.

### Study subjects

Healthy Taiwanese volunteers with a normal self-rated taste function enrolled this study. We excluded those with a history of oral or middle ear surgery, or acute oral infections. All eligible subjects were separated into two gender groups (male and female), and then each gender group into three age groups: 20-39 years, 40-59 years, and ≥60 years. A total of 40 subjects were included in each age group. They each took a whole mouth suprathreshold taste test, along with a taste quad test to measure their taste functions.

### Taste tests

Two solution-based taste tests were used; the whole-mouth suprathreshold test and the taste quad test. Between these two tests, subjects were allowed a break of 10-minutes.

### Whole mouth suprathreshold taste test

Five different suprathreshold concentrations of 4 basic tastant solutions were used in the whole mouth suprathreshold taste test [15]. Specifically, they were prepared as follows. Powders of sucrose, citric acid, sodium chloride (I Chan chemical Ltd., Taipei, Taiwan), and caffeine (Uni-Onward Corp., New Taipei City, Taiwan) were individually dissolved in distilled water to prepare the following tastant solutions: (a) sweet solution (concentrations of sucrose: 0.08, 0.16, 0.32, 0.64, 1.28 molar), (b) sour solution (concentrations of citric acid: 0.0026, 0.0051, 0.0102, 0.0205, 0.0410 molar), (c) salty solution (concentrations of sodium chloride: 0.032, 0.064, 0.128, 0.256, 0.512 molar), and (d) bitter solution (concentrations of caffeine: 0.0026, 0.0051, 0.0102, 0.0205, 0.0410 molar).

Prior to the beginning of the test, each subject was instructed not to smoke or eat for at least one hour. A small cup containing 10 mL of each tastant solution was then presented to the subject in a counterbalanced order. The solution in the cup was sipped, swished in the mouth for 10 seconds, and expectorated. The subject was then asked to indicate which taste the solution was, and rated the intensity and unpleasantness/pleasantness of the solution on 9-point scale in a forced-choice paradigm (i.e., forced to choose which one was the correct tastant, even if in doubt). The intensity of the solution was rated using a 9-point scale: 1: not present at all, 2: very slight, 3: slight, 4: definitely present, 5: moderate, 6: moderately strong; 7: strong; 8: very strong; 9: extremely strong. The pleasantness of the solution was rated using a 9-point scale: 1: dislike extremely, 2: dislike very much, 3: dislike moderately, 4: dislike slightly, 5: neither like or dislike, 6: like slightly; 7: like moderately; 8: like very much, 9: like extremely. Between successive cups, the mouth of the subject was rinsed with distilled water. Each of the 5 suprathreshold concentrations for the 4 tastant solutions was tested twice by the subject. Therefore, a total of 40 tests (4 tastants × 5 concentrations × 2 trials) were performed to generate a maximum score of 40. The recording data sheet of the whole mouth taste test is shown in Fig 1.

**Fig 1.**
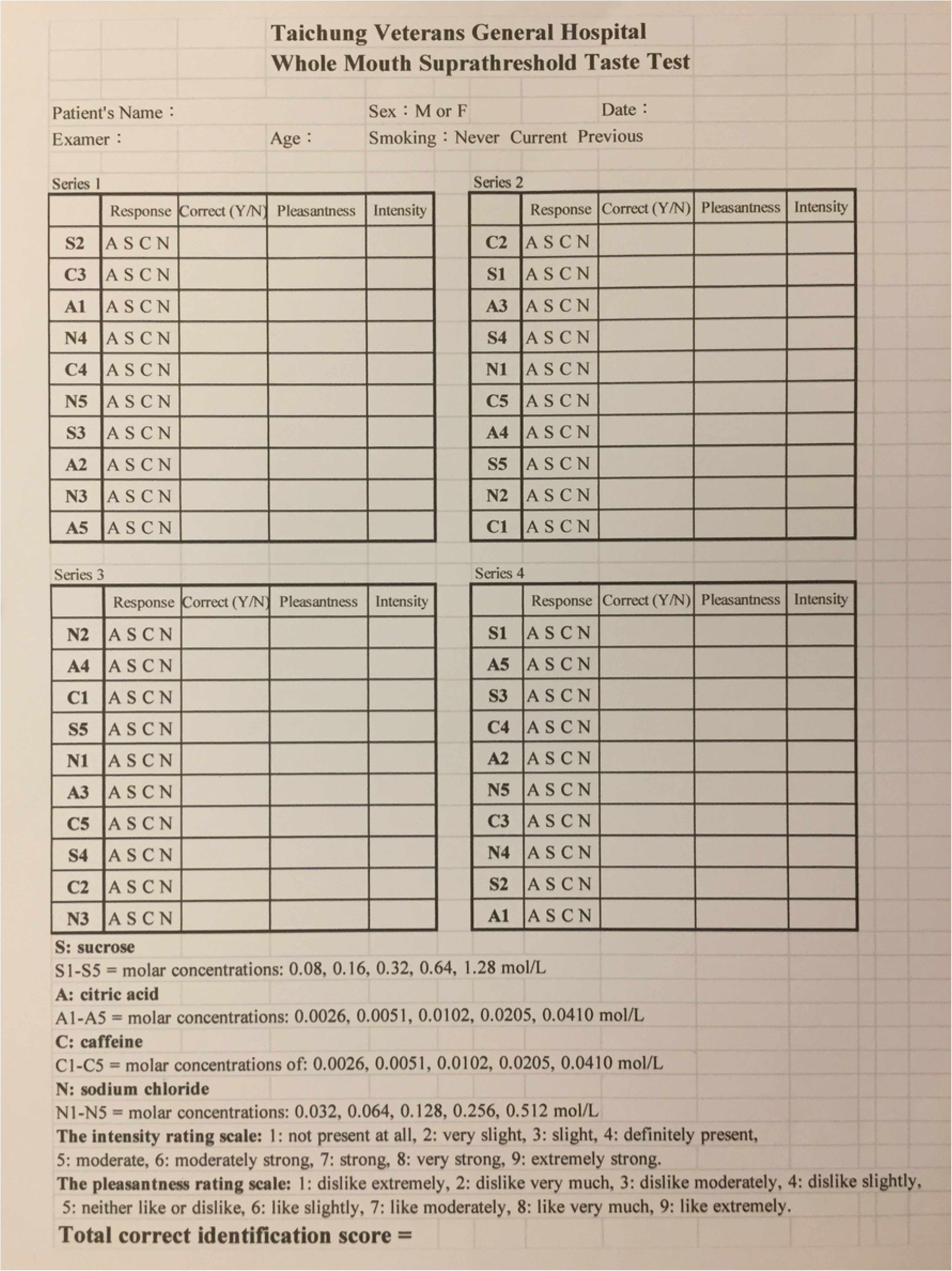
recording data sheet of the whole mouth taste test

### Taste quad test

Here, a single suprathreshold concentration of solution was prepared for each of the 4 basic tastants as follows: a 0.49 mol/L sucrose solution (for sweet), a 0.015 mol/L citric acid solution (for sour), a 0.31 mol/L sodium chloride solution (for salty), and a 0.04 molar caffeine solution (for bitter). Prior to beginning the test, the subject was instructed to protrude tongue, which was visually divided by the experimenter into 4 quadrants (quadrant 1: right posterior tongue, quadrant 2: right anterior tongue, quadrant 3: left anterior tongue, and quadrant 4: left posterior tongue). Next, using a micropipette, the experimenter dripped 15 μL of the tastant solution onto one of the 4 quadrants. In a forced-choice paradigm, the subject indicated which taste the solution was presented, and then rated the intensity of the solution on the same 9-point scale as that was used in the whole-mouth test. Then, the mouth was rinsed with distilled water. On each tongue quadrant, 4 tastant solutions were tested 6 times, with the tastants presented in a counterbalanced order. Therefore, a total of 96 tests were performed for each subject to generate a maximum score of 96. The recording data sheet of the taste quad test is shown in Fig 2.

**Fig 2.**
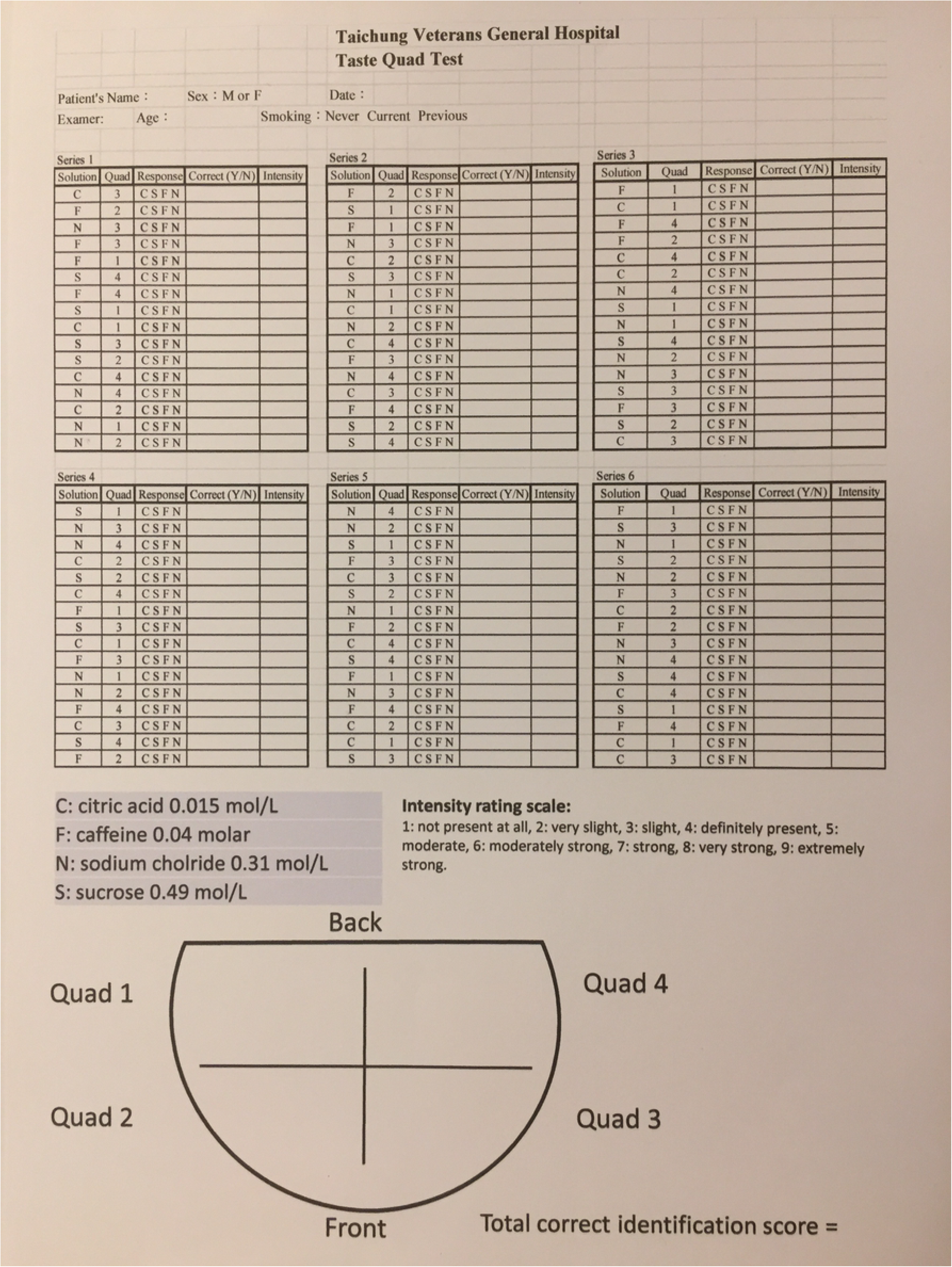
recording data sheet of taste quad test

### Statistical analyses

Descriptive data were presented as mean ± standard deviation. In both the whole mouth test and quad test, the correct quality identification scores, intensity and unpleasantness/pleasantness rating scores were compared across age groups using the Kruskal Wallis test, and compared between each two groups using the Dunn-Bonferroni test. The correct quality identification scores of the tastant solutions in each age group were then compared across the 4 tongue quadrants using the Friedman test. A score of correct quality identification at the 10 percentile was defined as the normative data [6,14]. All computations were performed using SPSS version 20 (Armonk, NY: IBM Corp.). Two-tailed p-values <0.05 were considered to be statistical significant.

## Results

### Subjects

We studied a total of 240 subjects whose age ranged from 20 to 88 years, with a mean of 48.04 years. Ages were not different between the male and female subjects for all 3 age groups (p= 0.086, 0.452, 0.369, respectively).

### Whole mouth suprathreshold taste test

The correct quality identification scores with the intensity and unpleasantness/pleasantness ratings of the tastant solutions are shown in Table 1.

**Table 1.**
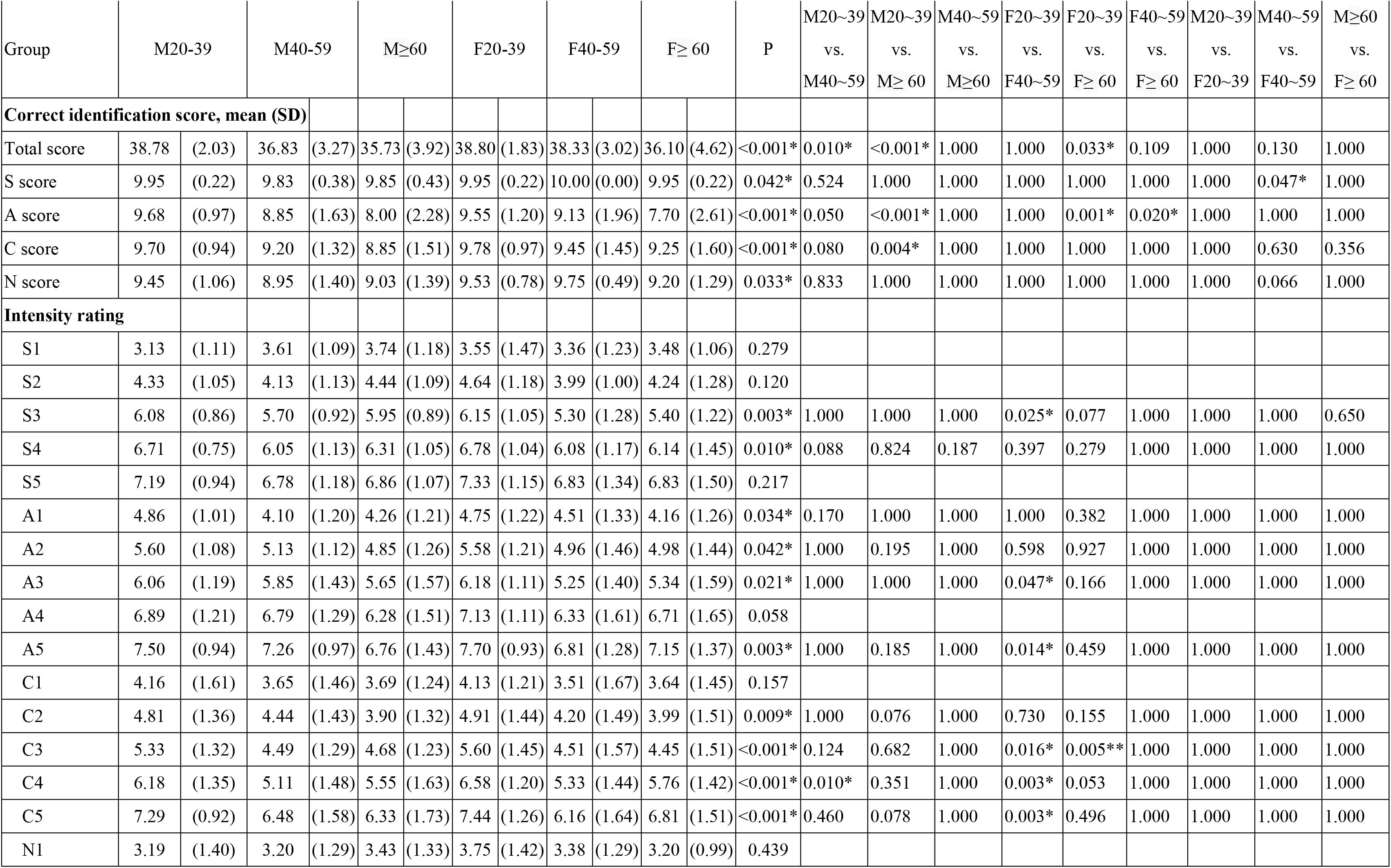
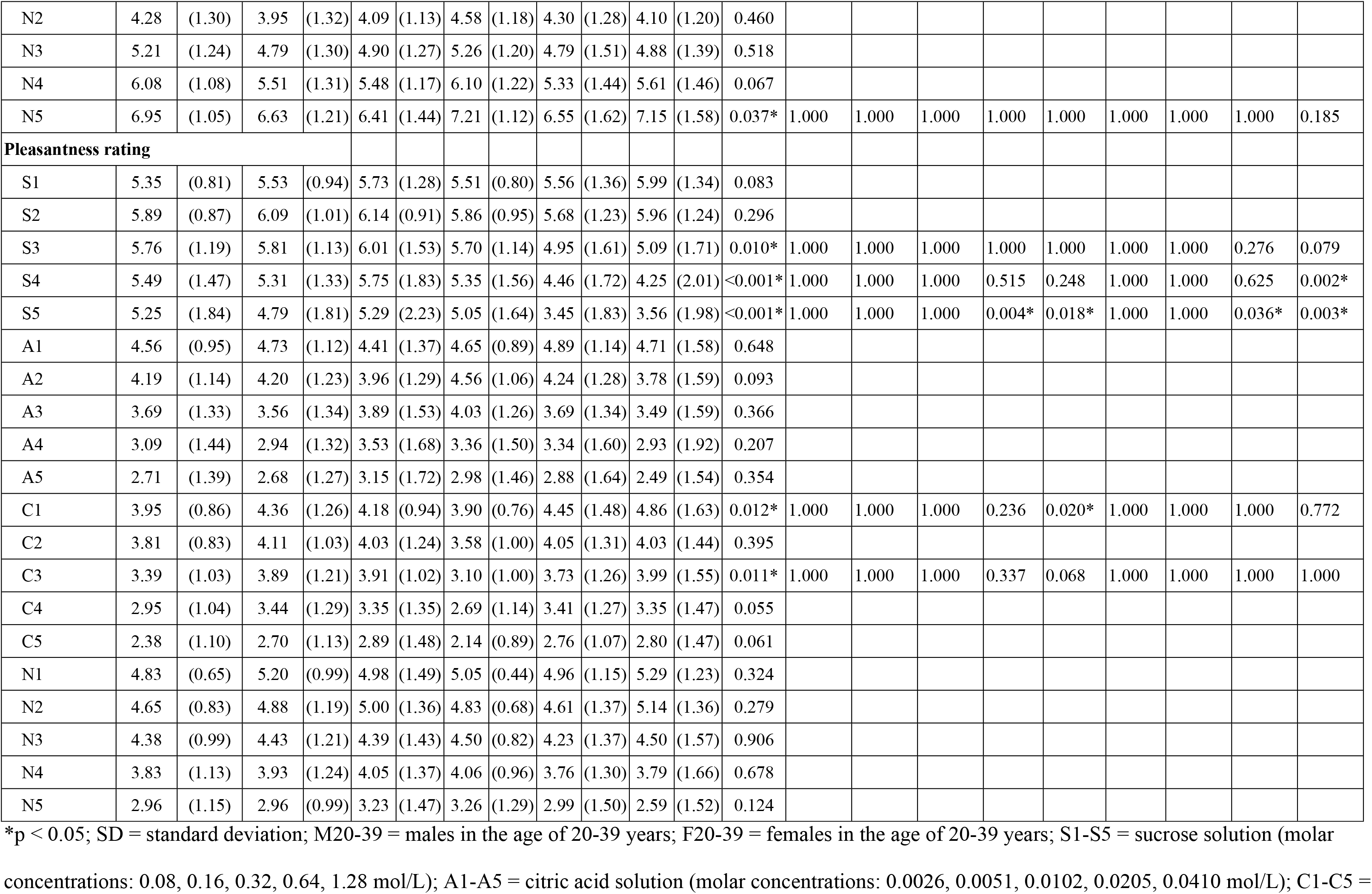
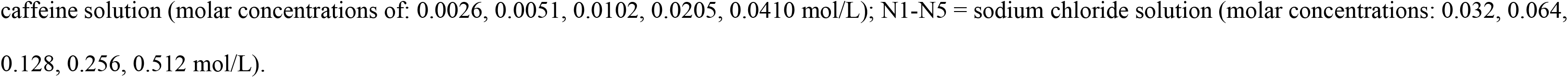
The whole mouth suprathreshold taste test.

The total scores of correct quality identification of the 4 tastant solutions was significantly higher for the age group of 20-39 years than the other two age groups regardless of gender ((vs 40-59 years: p= 0.01 for male, p=0.033 for female; ≥60 years: p <0.001 for male, p=0.033 for female). For individual tastants, male subjects in the age group 20-39 years had significantly higher scores for sour and bitter solutions than those ≥60 years (p= <0.001 for sour; p= 0.004 for bitter). Female subjects in age group ≥60 years had significantly lower scores for the sour solution than those younger in (20-39 years, p= 0.001; 40-59 years, p=0.02). Similar performances were found in both male and female subjects in the identification of sour, bitter and salty tastes. Nevertheless, females in the age group of 40-59 years had better identification for sweet than male subjects (p = 0.047).

The intensity rating scores were higher in the age group of 20-39 years, regardless of gender. In regard to taste preference, female subjects in the age group of 20-39 years liked sweet solution more than older female subjects. On the other hand, male subjects in the two older age groups liked sweet more than the female subjects (p= 0.036 for 40-59 years; p= 0.003 for ≥60 years).

### Taste quad test

Results (total and individual scores) of correct quality identification and intensity ratings of the 4 tongue quadrants are shown in Table 2.

**Table 2.**
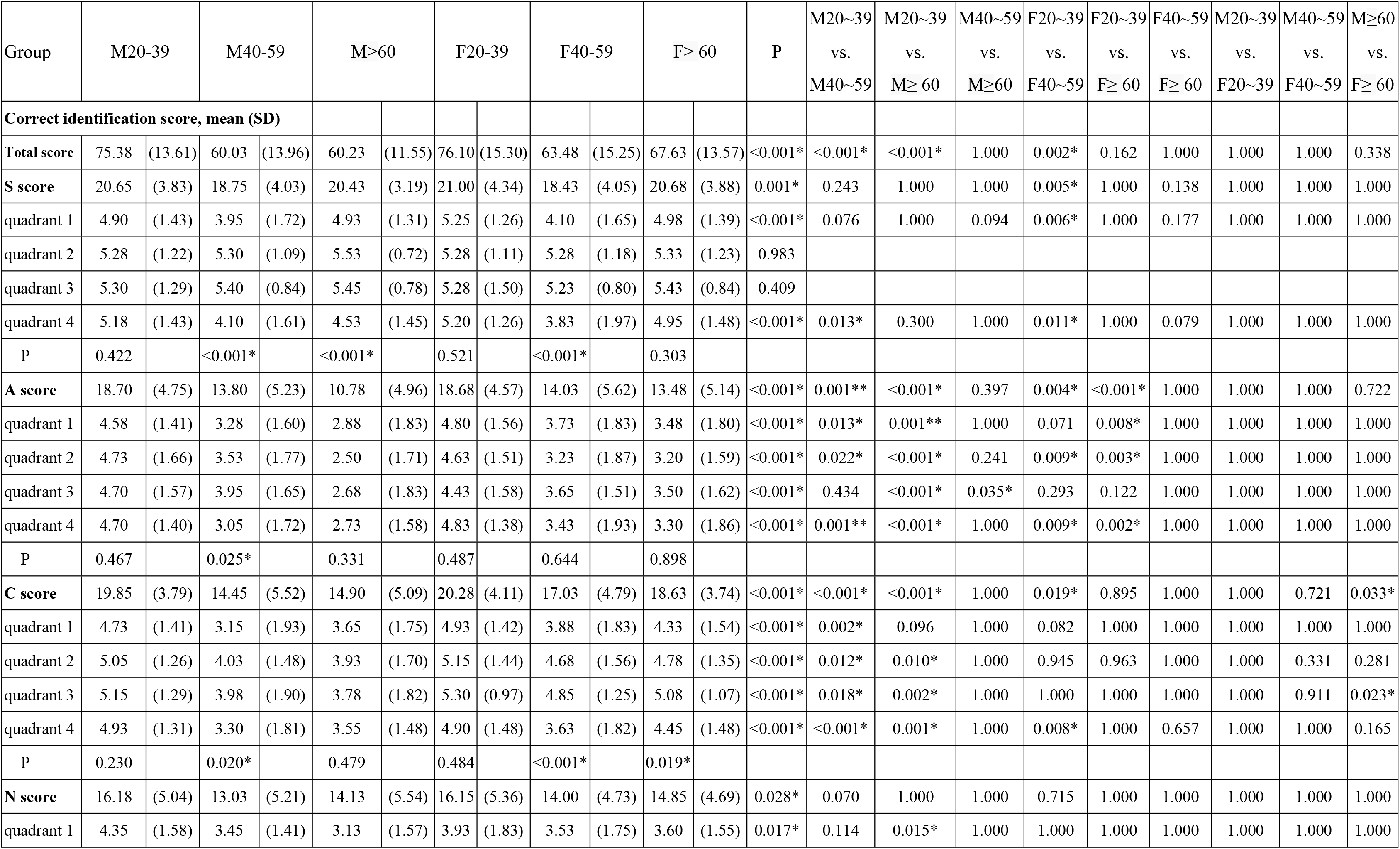
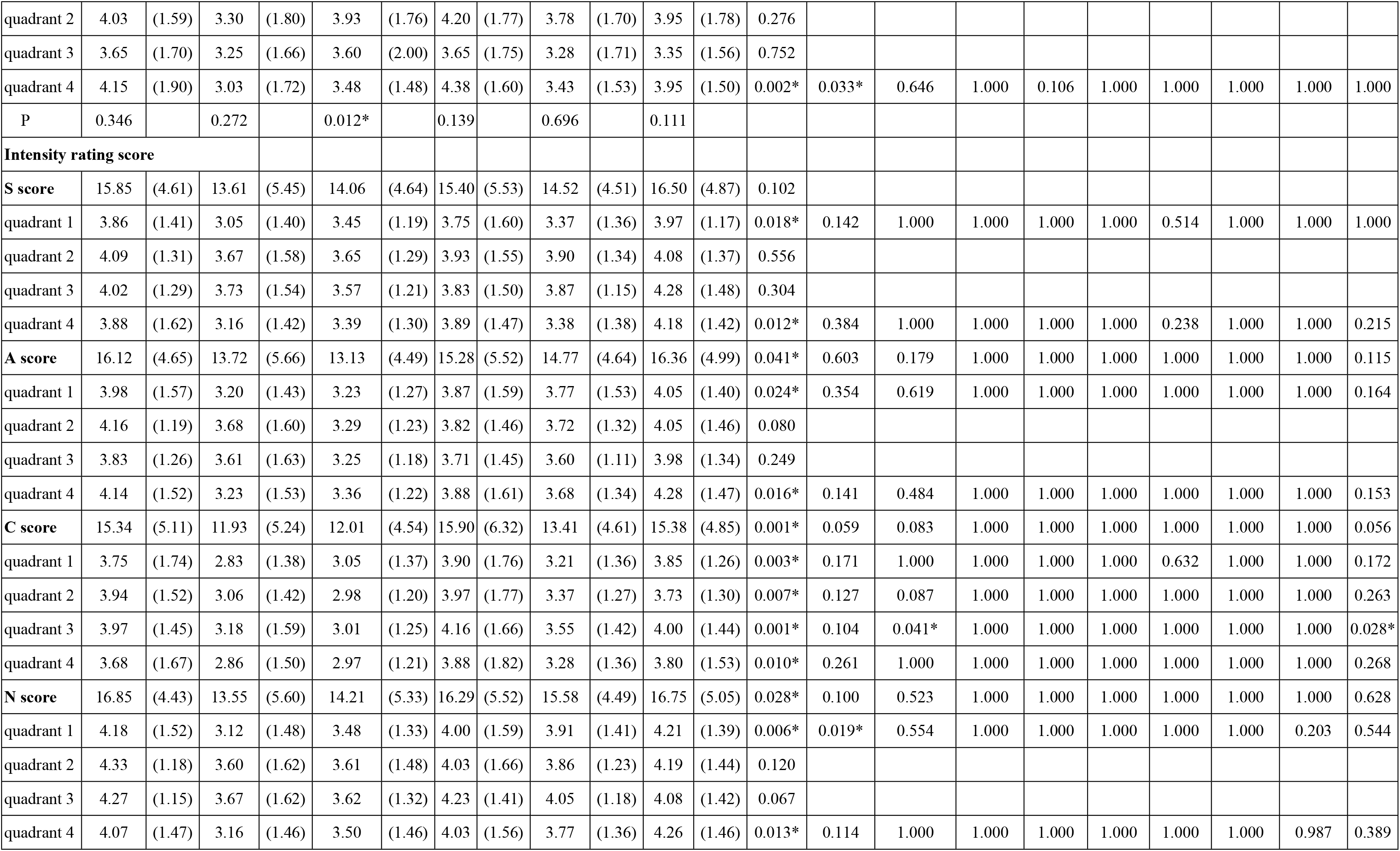
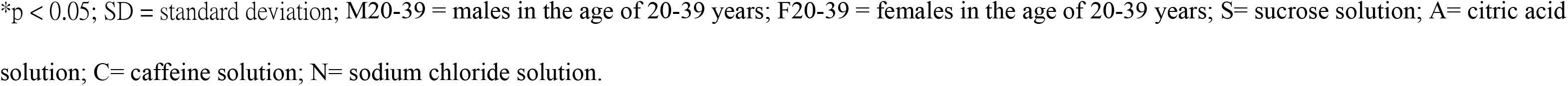
The test quad test.

The total scores for male subjects in the young age group (20-39 years) were significantly higher than the two older age groups (40-59 years, ≥60 years, both p <0.001). For female subjects, younger subjects had also higher scores than in the age group of 40-59 years (p= 0.002). In regard to individual quadrant, the correct identification scores appeared to worsen with age. A significant age influence of age was found in identifying sour and bitter tastants at the anterior tongue (quadrants 1 and 4), especially in male subjects. No significant gender differences were found in the total and individual scores of correct quality identification and intensity ratings except bitter tastant. Here male subjects ≥60 years had significantly lower identification and intensity rating scores than female subjects. When the correct quality identification scores were compared amongst the 4 tongue quadrants, we found no gender differences in the age group of 20-39 years. For those of 40-59 years, the correct quality identification scores of anterior tongue (quadrants 2and 3) were significantly higher than those of posterior tongue (quadrants 1and 4) for sweet, sour, and bitter tastant solutions in males aged 40-59 years, and for sweet and bitter tastants in females aged 40-59 years.

### Normative data

The normal data of the whole mouth suprathreshold taste test and test quad test are shown in Table 3. A score of correct quality identification at the 10th percentile was used to differentiate normogeusia to hypogeusia defined as the normative data [14].

**Table 3.** Normative data of the whole mouth suprathreshold taste test and test quad test.

| Group<br>(years of age) | Male<br>(20-39) | Male<br>(40-59) | Male<br>($\geq 60$) | Female<br>(20-39) | Female<br>(40-59) | Female<br>($\geq 60$) |
| --- | --- | --- | --- | --- | --- | --- |
| **Whole mouth suprathreshold taste test** |  |  |  |  |  |  |
| Minimum | 33 | 26 | 28 | 31 | 25 | 24 |
| Maximum | 40 | 40 | 40 | 40 | 40 | 40 |
| Tenth percentile | 35 | 32.1 | 29 | 37 | 34 | 26.3 |

| Taste quad test |  |  |  |  |  |  |
| --- | --- | --- | --- | --- | --- | --- |
| Minimum | 40 | 36 | 38 | 38 | 36 | 29 |
| Maximum | 91 | 86 | 86 | 96 | 89 | 90 |
| Tenth percentile | 55 | 42 | 46.2 | 53 | 42.3 | 48.2 |
| Quadrant 1 |  |  |  |  |  |  |
| Minimum | 6 | 4 | 5 | 7 | 6 | 8 |
| Maximum | 24 | 23 | 23 | 24 | 24 | 24 |
| Tenth percentile | 11 | 7.1 | 8.1 | 11.1 | 7.1 | 9.1 |
| Quadrant 2 |  |  |  |  |  |  |
| Minimum | 6 | 9 | 9 | 10 | 8 | 6 |
| Maximum | 24 | 24 | 22 | 24 | 22 | 23 |
| Tenth percentile | 13.2 | 12 | 11 | 12 | 10.1 | 11 |
| Quadrant 3 |  |  |  |  |  |  |
| Minimum | 8 | 11 | 9 | 8 | 11 | 10 |
| Maximum | 24 | 24 | 22 | 24 | 22 | 23 |
| Tenth percentile | 13.1 | 12.1 | 12 | 12 | 12.1 | 13 |
| Quadrant 4 |  |  |  |  |  |  |
| Minimum | 6 | 6 | 5 | 9 | 5 | 4 |
| Maximum | 24 | 23 | 22 | 24 | 24 | 24 |
| Tenth percentile | 11.2 | 7 | 8.1 | 12 | 8 | 12.1 |

## Discussion

Literature showed that the effect of aging on taste function is a decline during the aging process, and that this decline with age is probably tastant-dependent [4]. In our present study, we found similar decline in healthy Taiwanese subjects in terms of the total correct identification scores obtained from the whole mouth taste tests. Our findings are in discrepancy with those of Yang et. al., [8] who found no age-differences in scores obtained from the triple drop tests in healthy Chinese.

Welge-Lussen et. al., [13] found that women are slightly better than men in identifying different tastes. Doty et. al., [16] employed procedures like ours, and found no gender differences in their whole mouth taste test. Our study showed no gender differences in the same age group.

Regarding individual taste quality, sweet taste was reported to be more robust against age effects [13]. Our results from the whole mouth taste test further showed that in addition to sweet, the ability to identify salty quality was also age-independent. We found no gender differences to aging effect on individual taste, except women of 40-59 years had better identification scores for sweet than men of the same age group.

Regarding intensity and pleasantness rating in whole mouth test, we found female subjects of 20-39 years liked the sweet taste more than those in older ages. In addition, male subjects significantly liked sweet taste more than female subjects of same ages. It was reported that young subjects rate only bitter and sour stimuli more intensely than older subjects, but not sweet and salty stimuli [7]. Consistent with this, we found similar results in our whole mouth taste test, particularly for the bitter stimulus. But our subjects of 20-39 years also rated sweet stimulus more intensely than older subjects. When gender was reported to affect the judgment of taste intensity [7], our subject, regardless of gender, rated all individual stimuli with similar judgments.

The regional taste test is used to detect gustatory blind regions on the tongue surface [14]. Few studies reported that age-related taste declines on localized regions of the tongue [17]. Pingel et al., [14] found that the scores of the regional taste test worsen with aging, women score higher than men, and the left tongue surface is more sensitive than the right in elderly people. On the other hand, Doty et. al., [17] reported t a decline in age-related taste function at the front surface of the tongue occurs slightly during middle age, becoming marked as people grow older; without gender differences. In our study, the correct identification scores were similar across the 4 quadrants for subjects of 20-39 years, regardless of gender. Nevertheless, we found a decline in age-related taste function on the posterior tongue surface, especially for sweet and bitter tastes. Regarding gender differences, women had correct identification scores for bitter quality higher than men, particularly for older subjects (≥60 years).

Taste function is rarely investigated in Asian populations. It has been assumed that eating-habits, diet and cultural differences between populations affect the results of taste tests [8,18,19]. Yong et. al., [8] proposed that Asians are more taste-sensitive. Shu-Fen et. al., [19] reported that Indians have recognition thresholds for all taste qualities higher than Chinese. Our results came from a Taiwanese population base which exhibits eating-habits, diet and culture different from Western populations, as well as Indians and other Chinese populations. Future cross-cultural studies are needed to clarify such population-related differences.

In summary, our results showed that both age and gender affect the taste function in healthy Taiwanese. The total correct identification scores of the whole mouth test and test quad taste test decreased with age, but the influences are not uniform. For instance, sweet and salty tastes were robust to aging, and the age-related taste decline occurred mostly on the posterior tongue surface. Moreover, ethnicity of subjects may also influence taste perception. Therefore, in measuring taste functions and interpreting the results, multiple factors like age, gender, tongue region, taste quality and ethnicity should all be taken into consideration.

## Acknowledgements

The authors are grateful to the Biostatistics Task Force, Taichung Veterans General Hospital, Taichung, Taiwan, for their statistics assistance.

